## Supplementary material for "Social learning mechanisms shape transmission pathways through replicate local social networks of wild birds"

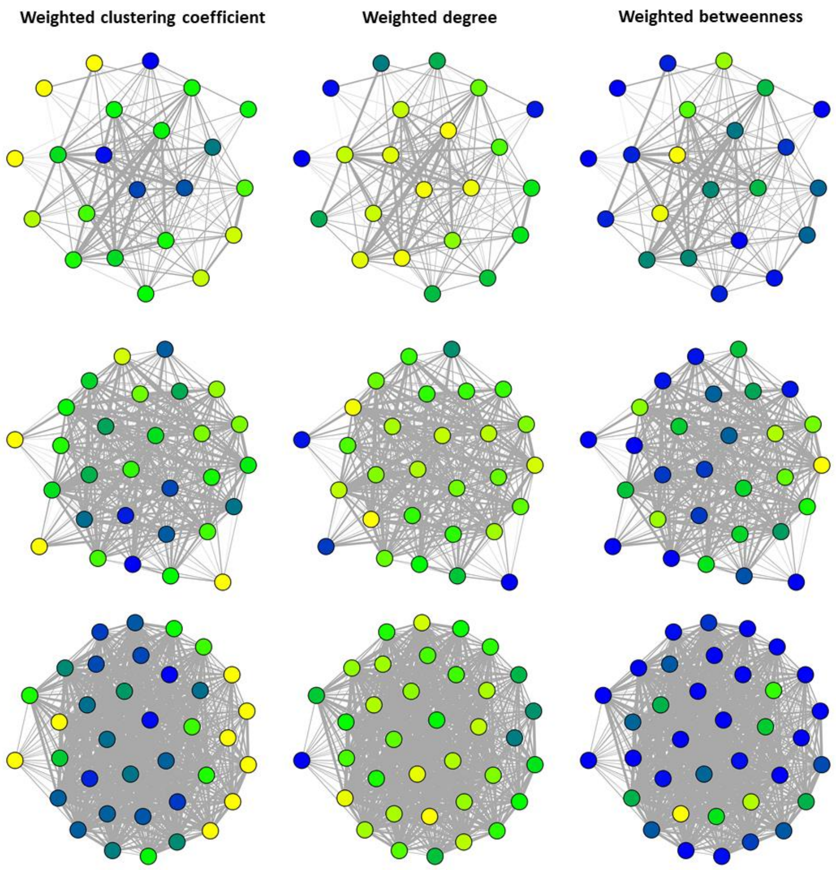


**Figure S1. Great tit social networks illustrating individuals' different social network positions.** Each row shows one weekly, local network of different size (top to bottom: 22, 29, 39 individuals). Columns show the three social network metrics (weighted clustering coefficient, weighted degree and weighted betweenness) and colours represent the range in network metrics (yellow indicates larger values and blue smaller values.

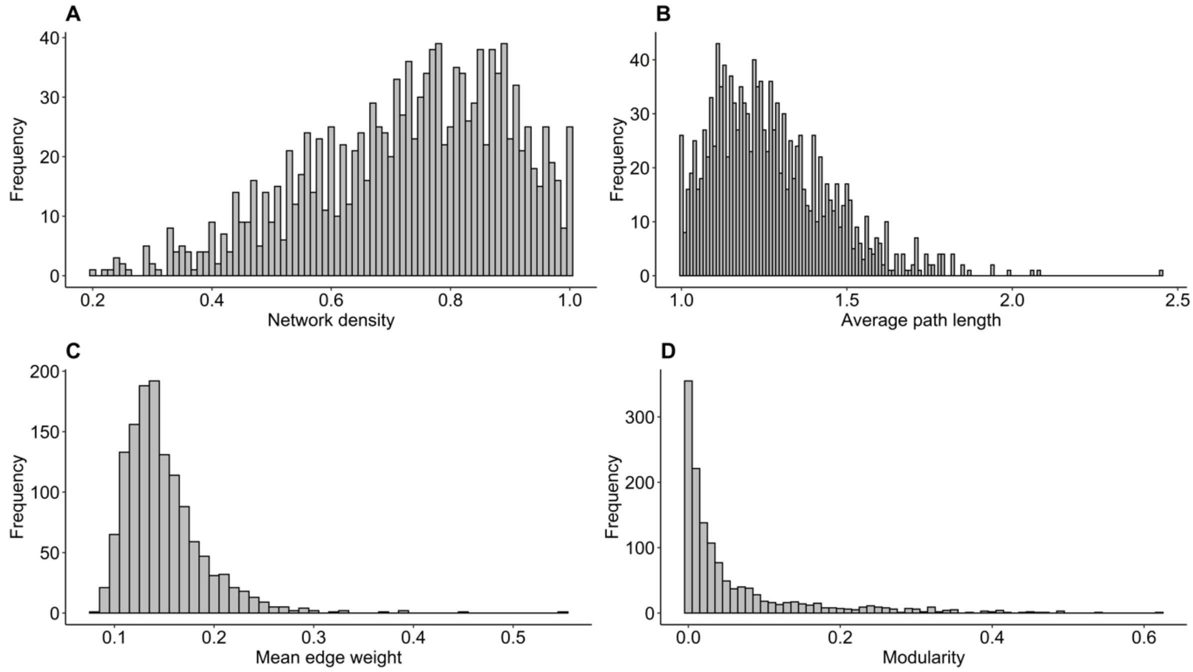


**Figure S2. Data distribution of four global network metrics from the 1343 weekly, local social networks. A**: Network density, calculated as the number of existing connections divided by all potential connections. **B**: Average path length, **C**: Mean edge weight defined as the average of all edge weights in a network (excluding zero edges). **D**: Modularity, inferred as the structural communities (modularity index Q) for each network using the edge betweenness community detection algorithm. We calculated the four global network metrics using the package 'igraph' (Csardi and Nepusz 2006). The network density was calculated as the number of existing connections divided by all potential connections. The global clustering coefficient was defined as the ratio of the triangles and the connected triples within the network. We inferred the structural communities (modularity index Q) for each network using the edge betweenness community detection algorithm. The average edge strength was defined as the average of all edge weights in a network (excluding zero edges).

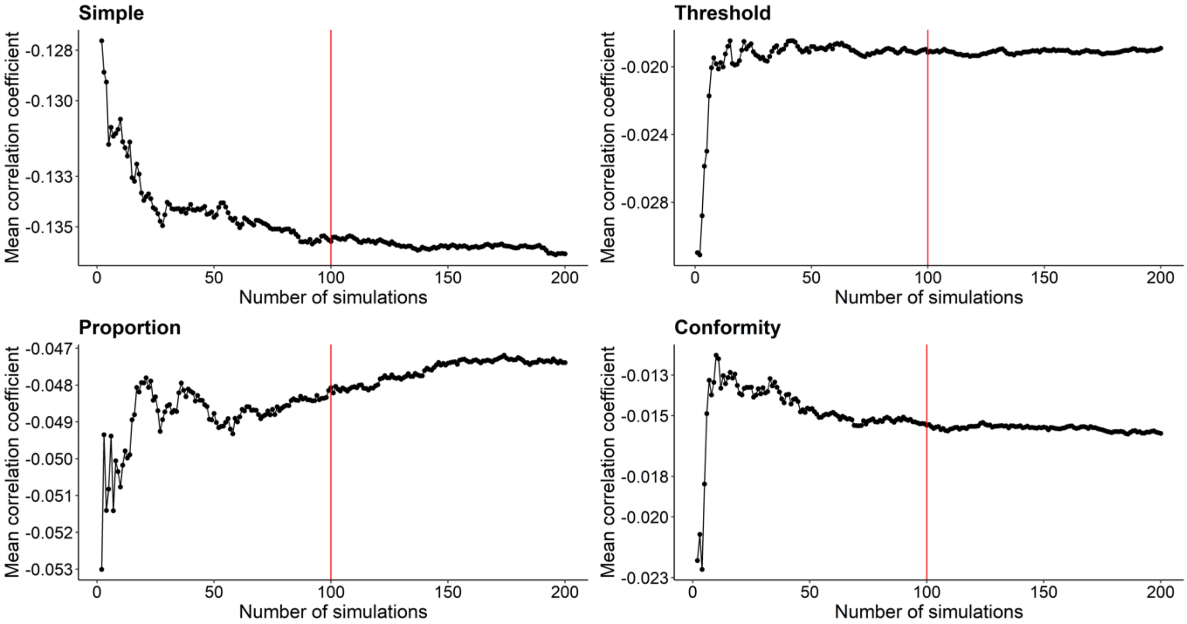


**Figure S3.** Mean correlation coefficient between weighted degree and order of acquisition in relation to different numbers of simulations. After about 100 simulation runs (red vertical line), mean correlation coefficients do not fluctuate anymore to a large extent. Therefore, 100 simulation runs were chosen as a meaningful cut-off.

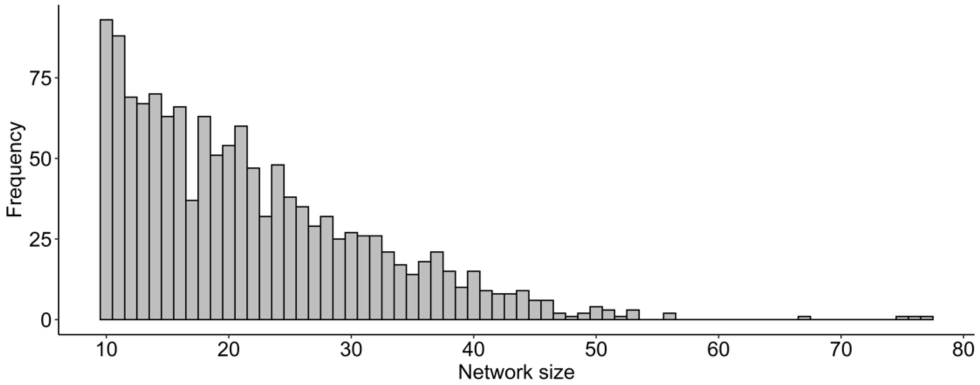


**Figure S4**. Data distribution of network sizes (i.e. the number of individuals each network contained).

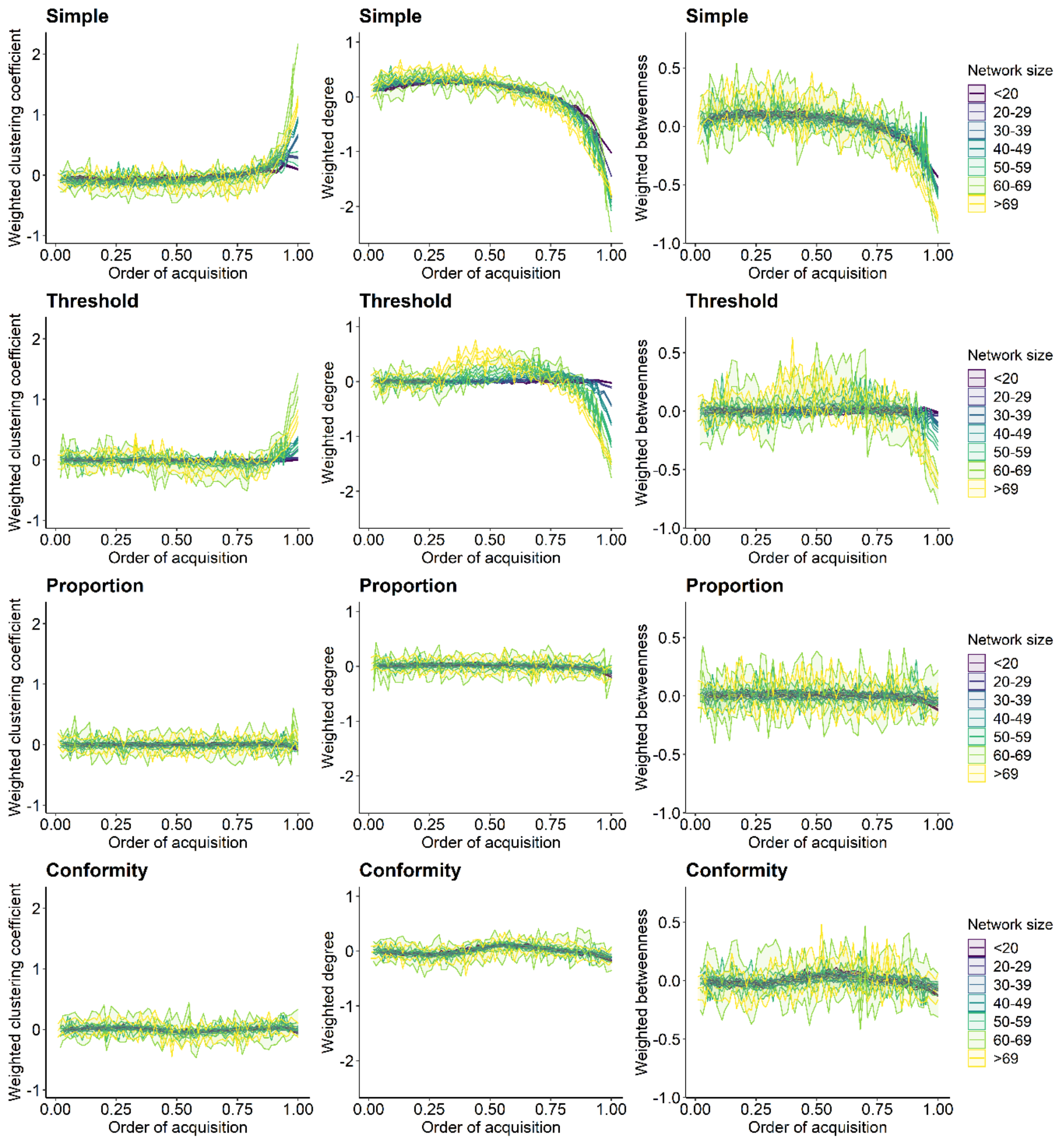


**Figure S5.** Relationship between individual network metric and the standardized order of acquisition (OAC) for each transmission rule and split by network size. Each column shows a different network metric (left to right: weighted clustering coefficient, weighted degree and weighted betweenness). Each row represents one of the four spreading rules (top to bottom: simple, threshold, proportion and conformity). A value of OAC=0.5 corresponds to 50% of individuals within the social network being knowledgeable. Lines show the average network metric for each OAC and ribbons show the 95% Confidence Interval from the 100 simulations for each binned group of network sizes. Colour represents network size with darker colour indicating smaller sizes. The social transmission rate ‘s’, the threshold location ‘a’, and the frequency-dependence ‘f’ were set to 5.

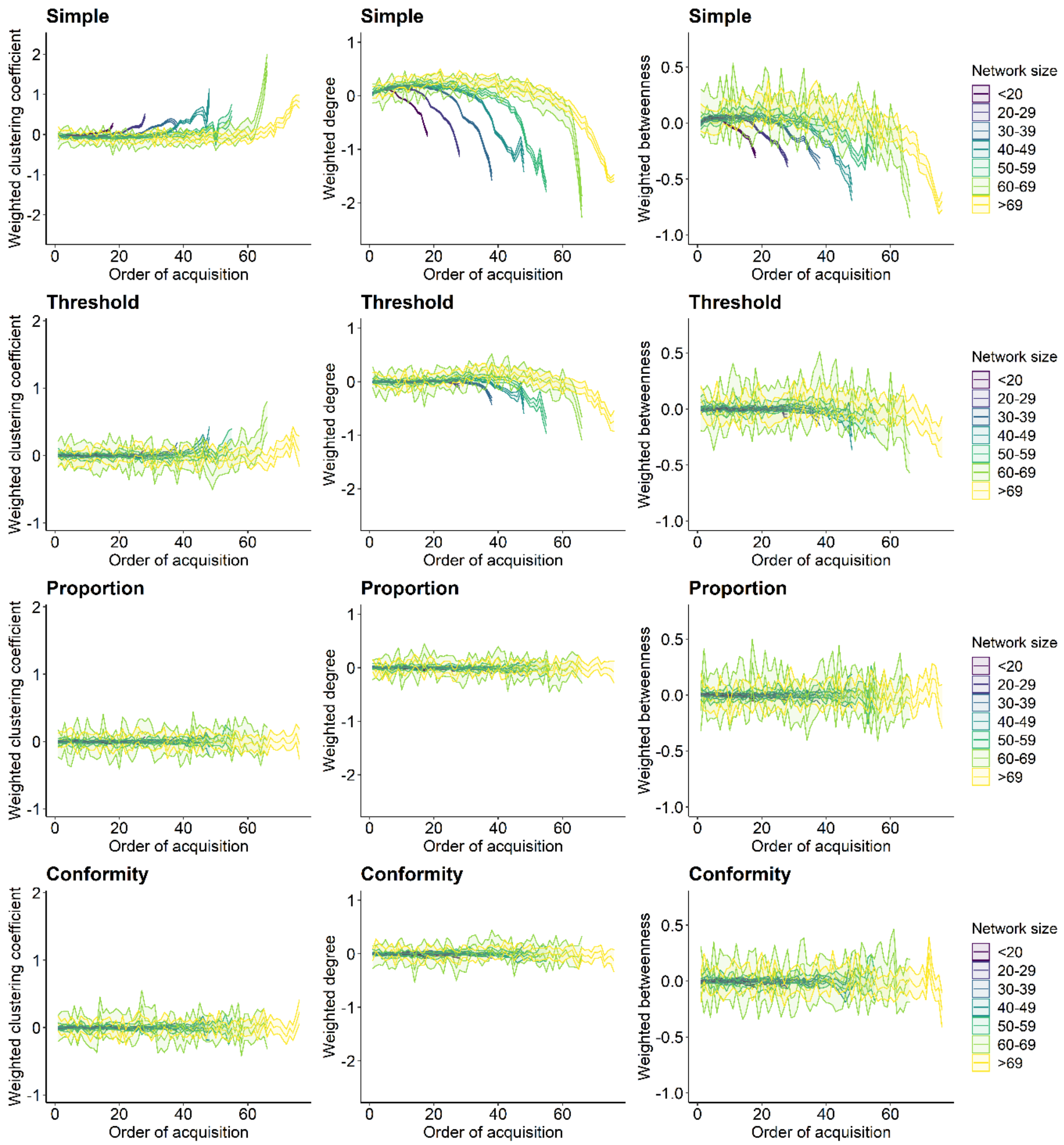


**Figure S6. Relationship between individual network metric and the order of acquisition for each social learning rule with a social learning rate of 1**. Each column shows a different network metric (left to right: weighted clustering coefficient, weighted degree and weighted betweenness). Each row represents one of the four spreading rules (top to bottom: simple, threshold, proportion and conformity). Lines plot the average network metric for each order of acquisition and ribbons show the 95% Confidence Interval from the 100 simulations for each binned group of and network sizes. Colour represents network size with darker colour indicating smaller networks. The threshold location ‘a’ and the frequency-dependence ‘f’ were set to 5.

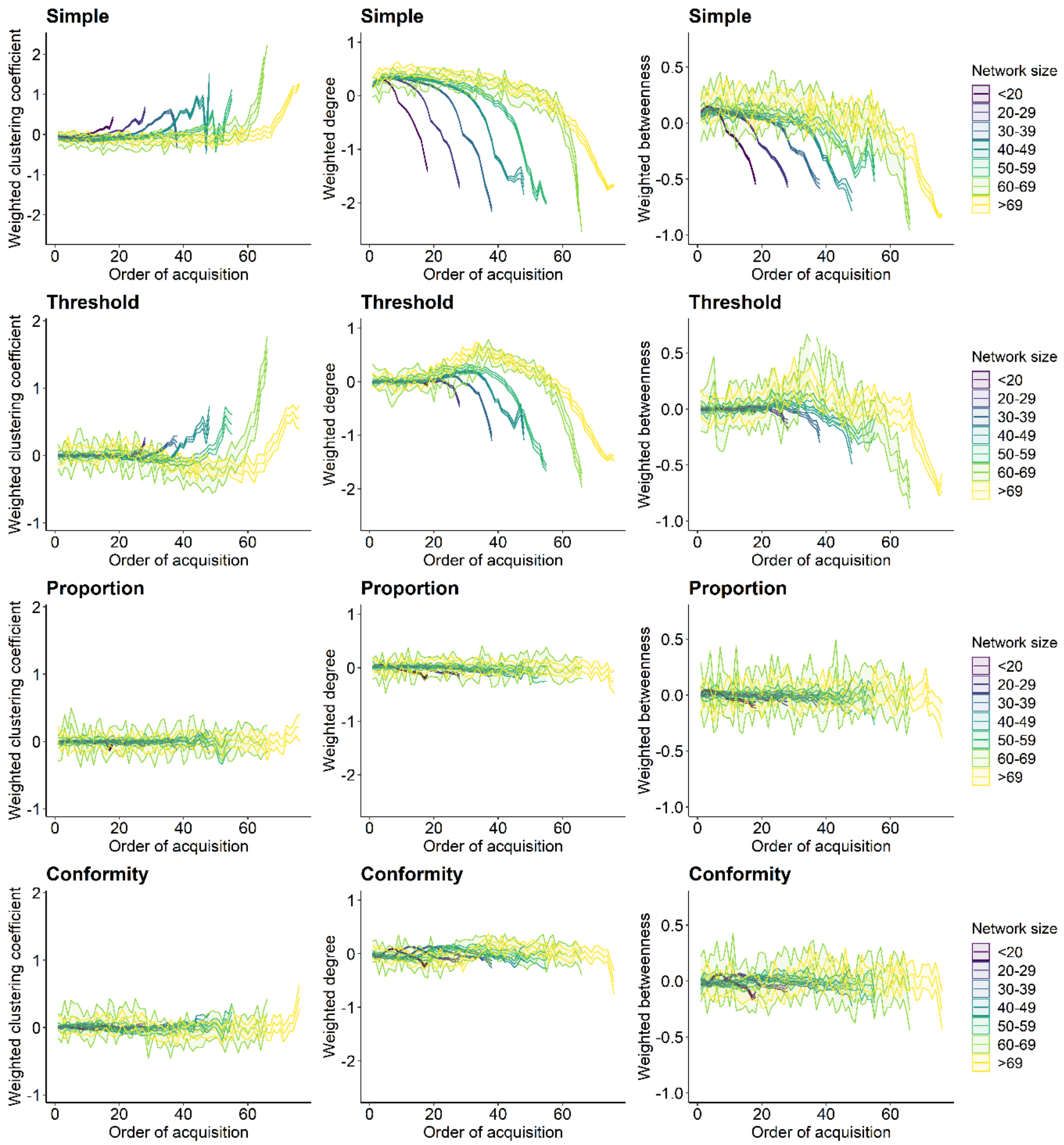


**Figure S7. Relationship between individual network metric and the order of acquisition for each social learning rule with a social learning rate of 10**. Each column shows a different network metric (left to right: weighted clustering coefficient, weighted degree and weighted betweenness). Each row represents one of the four spreading rules (top to bottom: simple, threshold, proportion and conformity). Lines plot the average network metric for each order of acquisition and ribbons show the 95% Confidence Interval from the 100 simulations for each binned group of and network sizes. Colour represents network size with darker colour indicating smaller networks. The threshold location ‘a’ and the frequency-dependence ‘f’ were set to 5.

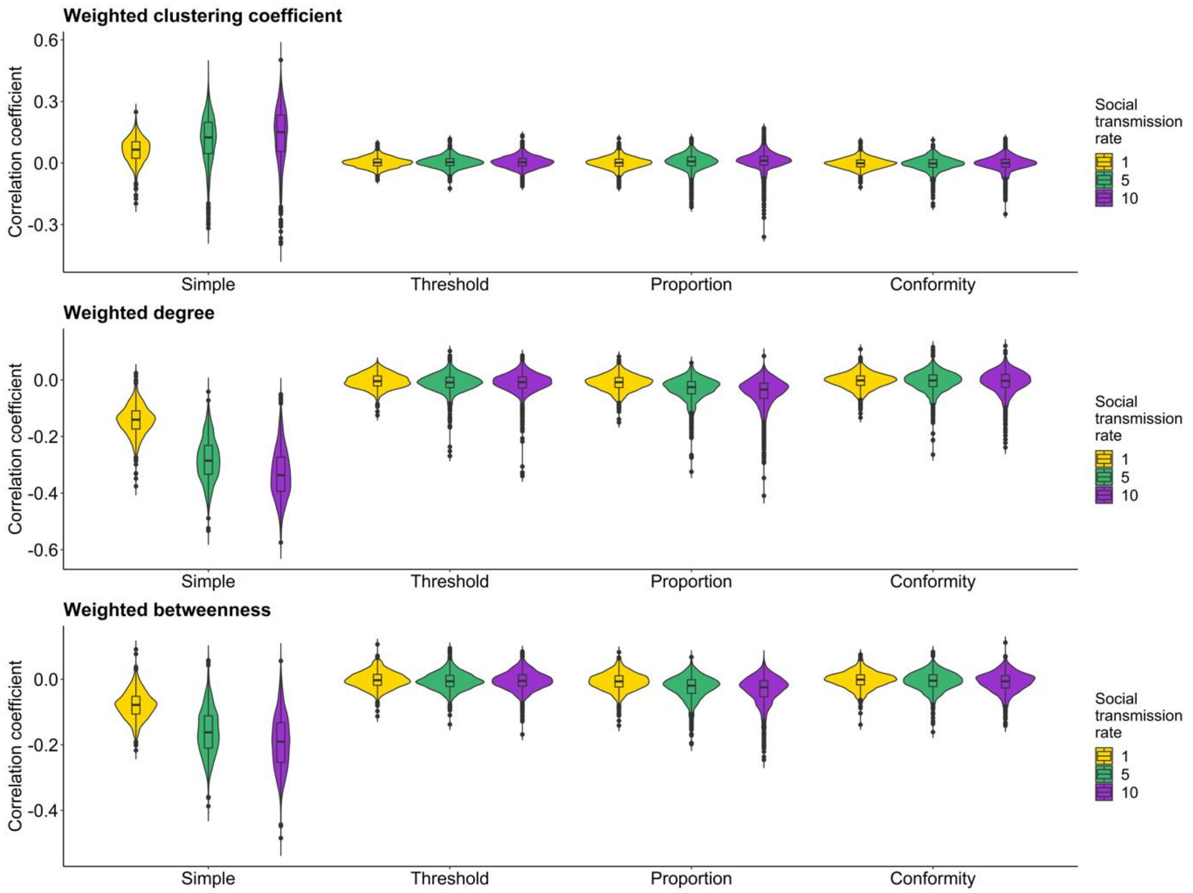


**Figure S8. Distribution of average correlation coefficients for each social learning rule and network metric under different social learning rates.** Violin and boxplots show the distribution of the average correlation coefficients across 100 simulations from each network for each of the four social learning rules (i.e. simple, proportion, conformity and threshold). Each plot shows one of the individual network metrics (weighted clustering coefficient, weighted degree, weighted betweenness). Colour represents the parameter set for the social transmission rate.

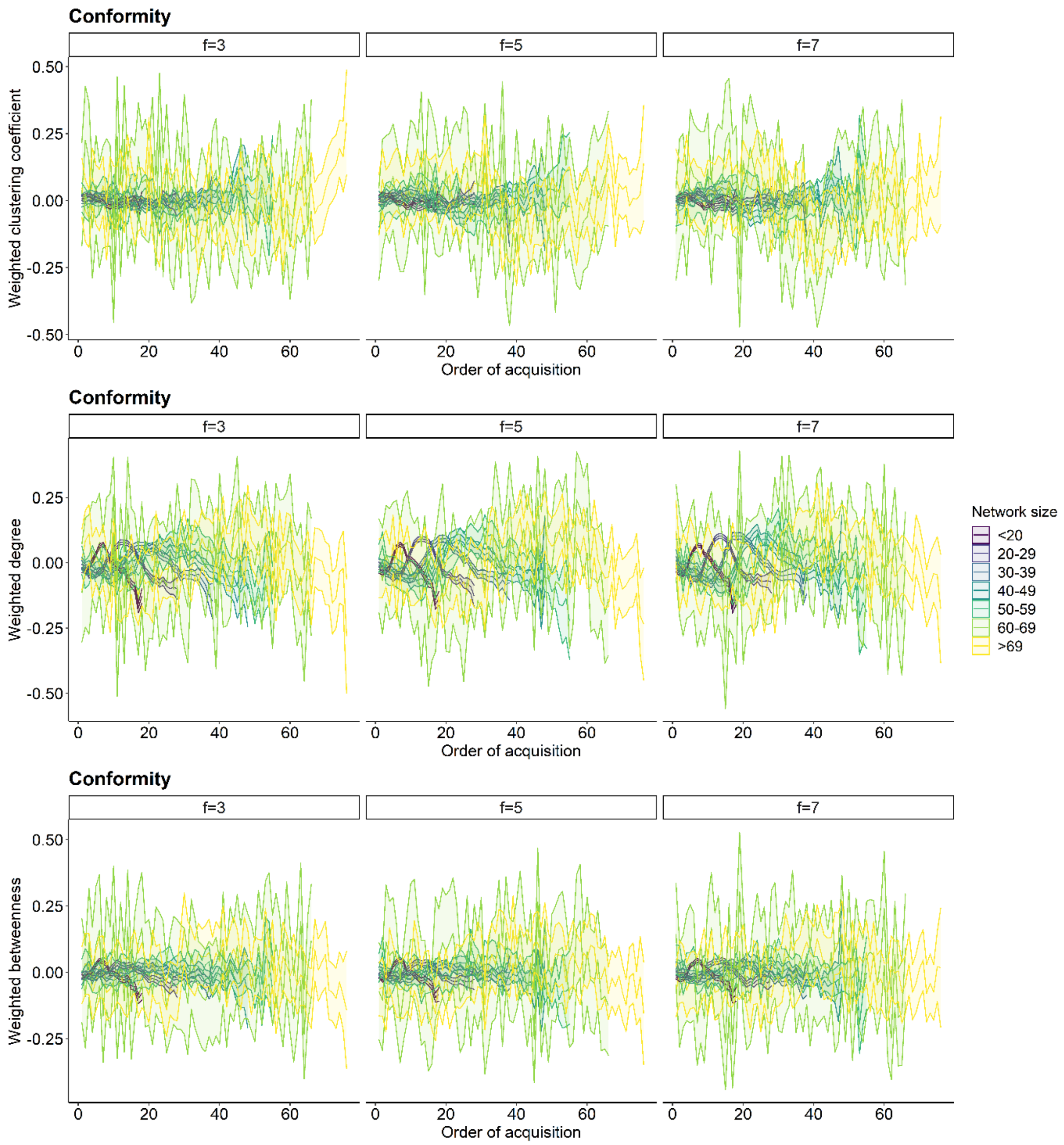


**Figure S9. Relationship between individual network metric and the order of acquisition for the conformity learning rule under different frequency-dependent values 'f'**. Each row shows a different network metric (top to bottom: weighted clustering coefficient, weighted degree and weighted betweenness). Each column represents a different parameter for the frequency dependence variable 'f'. Lines plot the average network metric for each order of acquisition and ribbons show the 95% Confidence Interval from the 100 simulations for each binned group of network sizes. Colour represents network size with darker colour indicating smaller networks. The social transmission rate was set to 5.

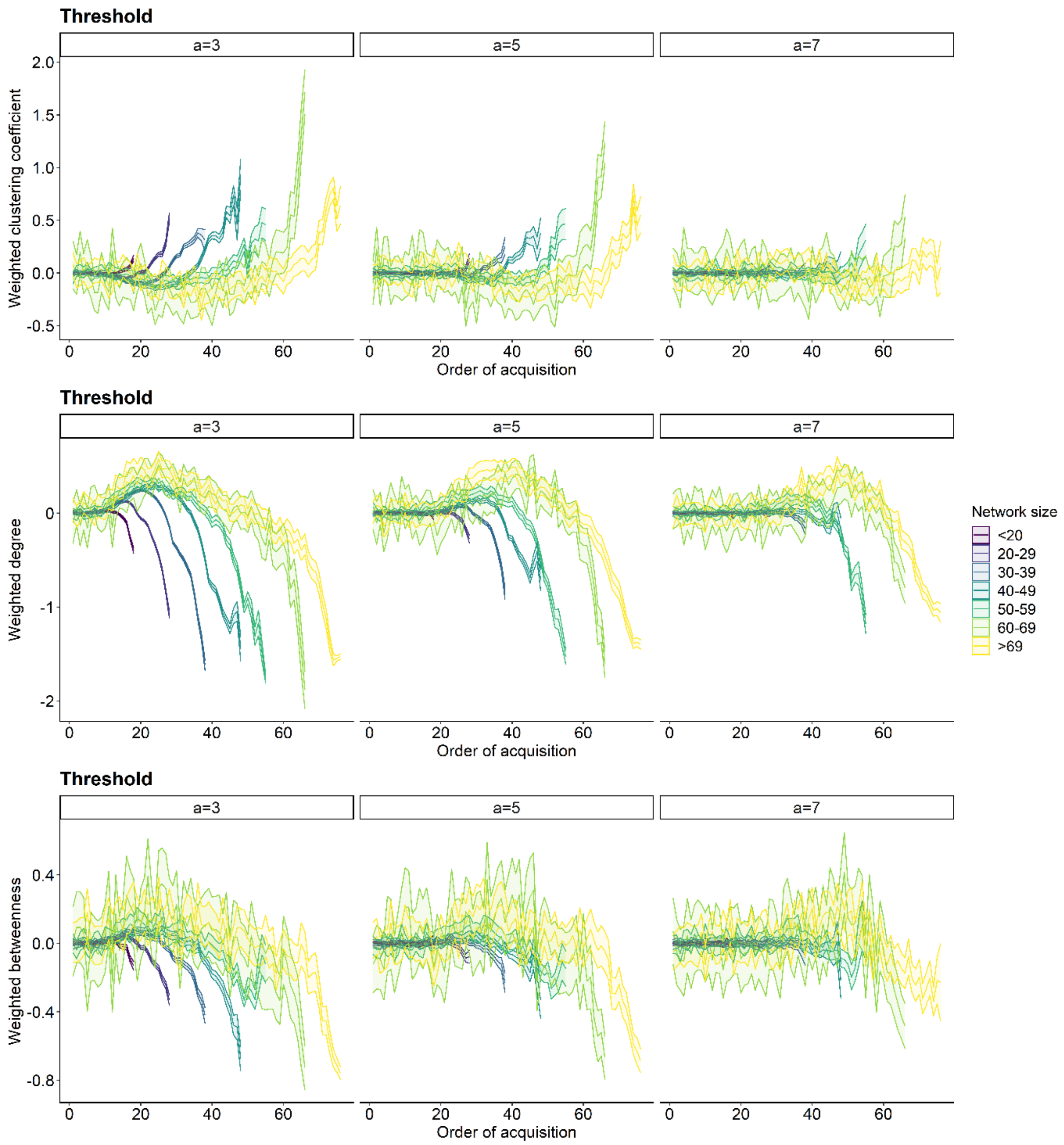


**Figure S10. Relationship between individual network metric and the order of acquisition for the threshold learning rule under different threshold locations ‘a’**. Each row shows a different network metric (top to bottom: weighted clustering coefficient, weighted degree and weighted betweenness). Each column represents a different parameter for the threshold location ‘a’ (3-7). Lines plot the average network metric for each order of acquisition and ribbons show the 95% Confidence Interval from the 100 simulations for each binned group of network sizes. Colour represents network size with darker colour indicating smaller networks. The social transmission rate was set to 5.

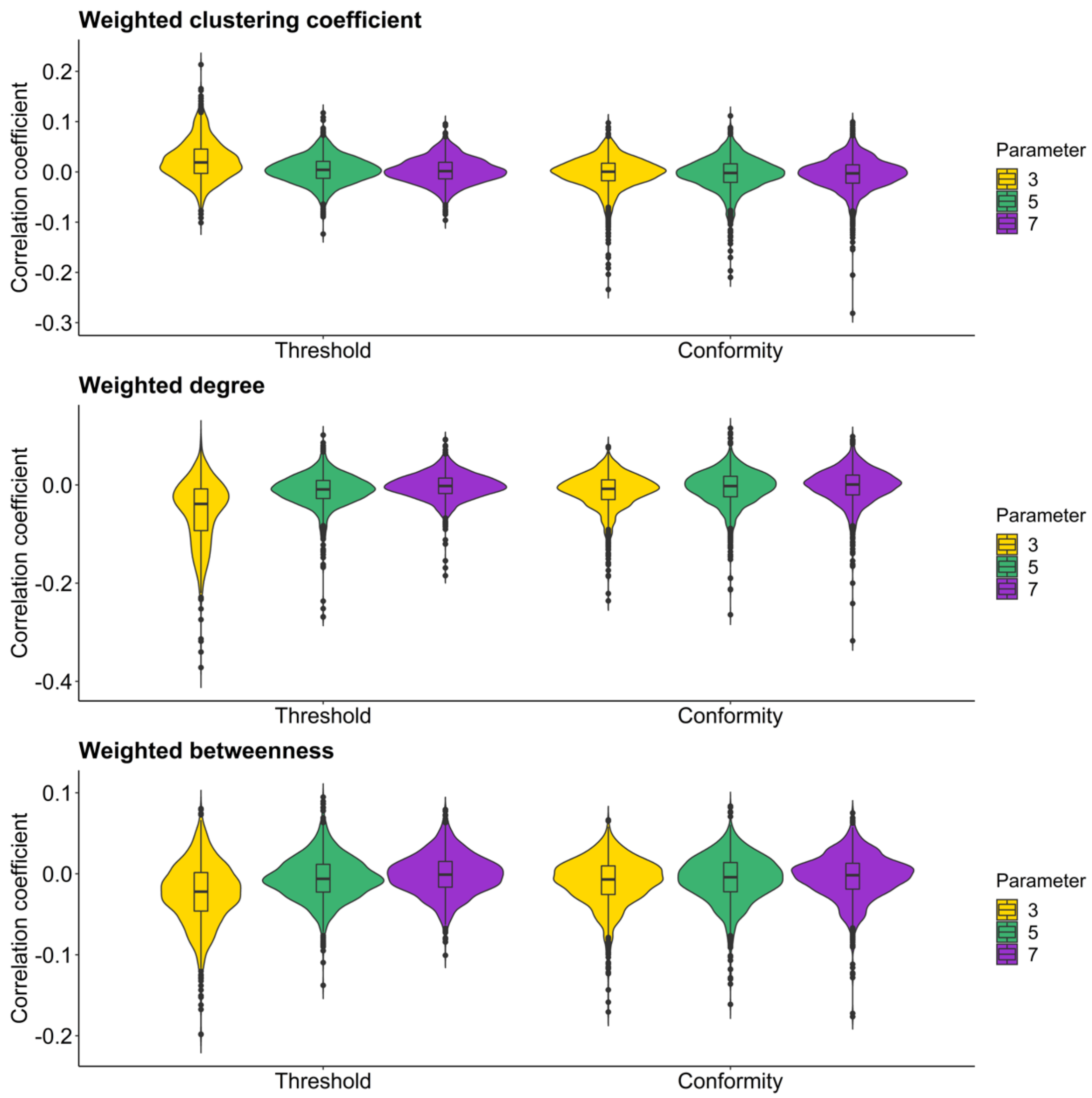


**Figure S11 Distribution of average correlation coefficients for the threshold and conformity learning rule under different parameters.** Violin and boxplots show the distribution of the average correlation coefficients across 100 simulations from each network for each of the four social learning rules (i.e. simple, proportion, conformity and threshold). Each plot shows one of the individual network metrics (weighted clustering coefficient, weighted degree, weighted betweenness). Colour represents the parameter set for the social transmission rate.

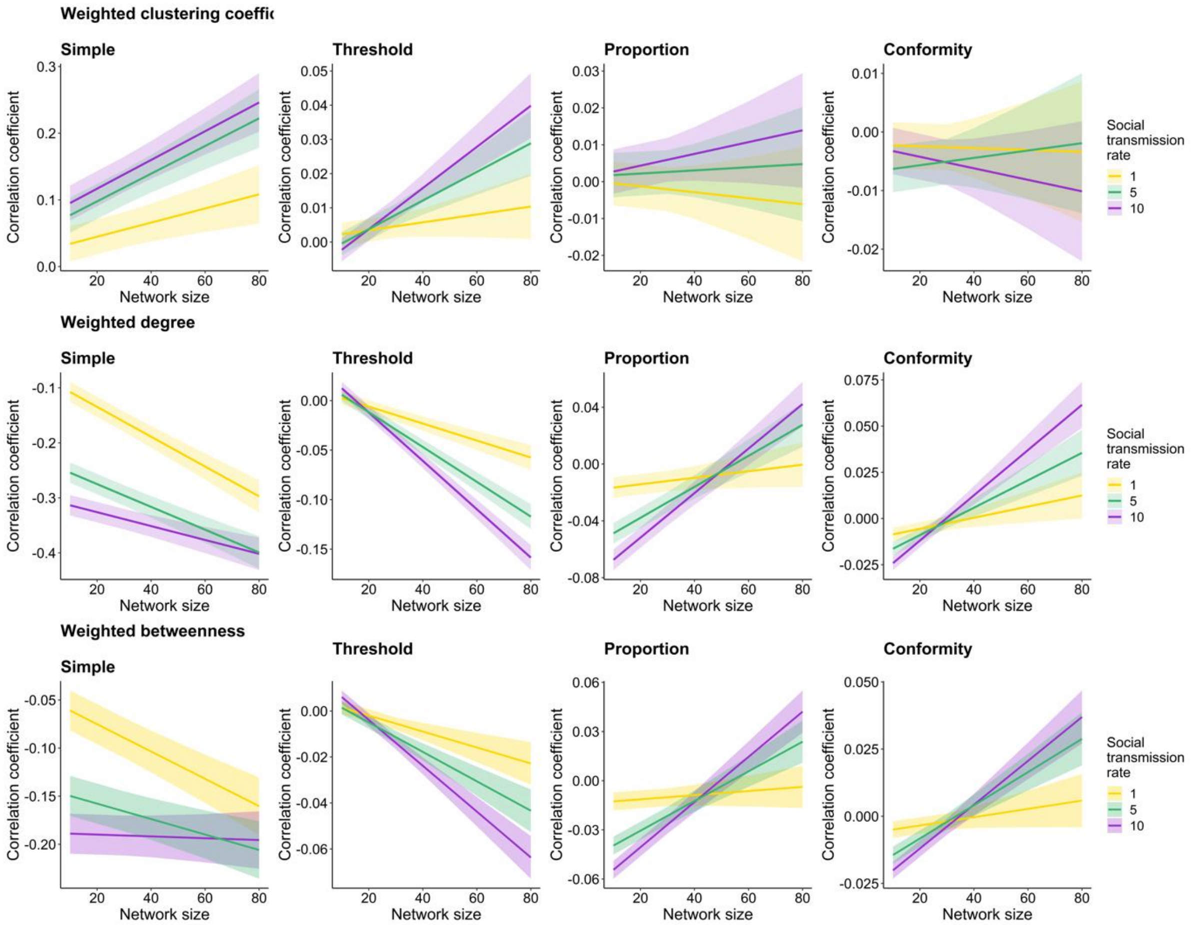


**Figure S12. Relationship between correlation coefficient and network size across the four social learning rules under different social learning rates.** Each row shows one of the individual network metrics (top to bottom: weighted clustering coefficient, weighted degree and weighted betweenness) and each column a different social learning rule (left to right: simple, threshold, proportion, conformity). Lines show the predicted effects generated from linear mixed-effect models (LMM) and ribbons show the 95% confidence intervals (see Table S1 for model results). Colour represents the different parameters set for the social transmission rate.

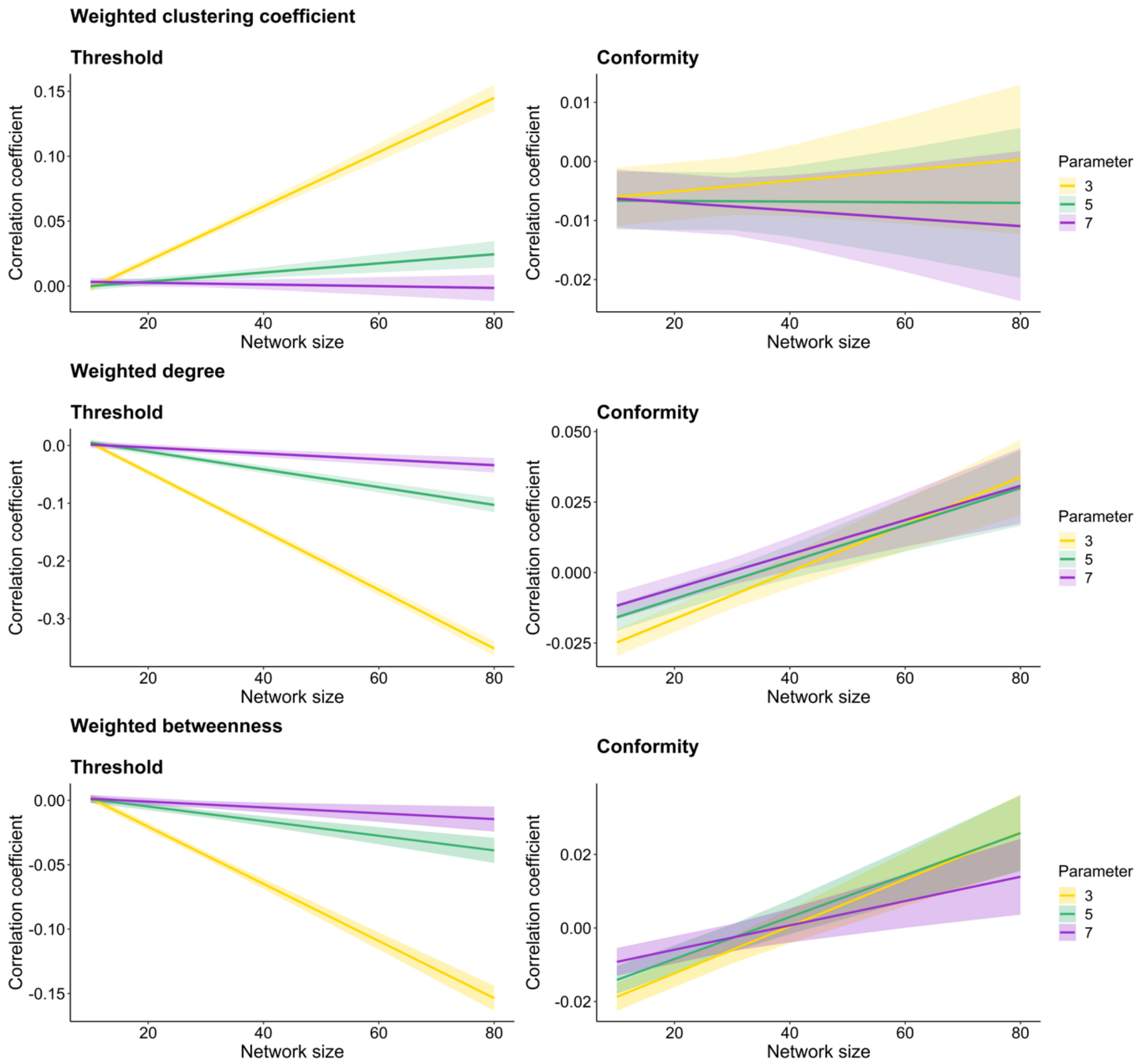


**Figure S13. Relationship between correlation coefficient and network size for the threshold and conformity learning rule under different parameters.** Each row shows one of the individual network metrics (top to bottom: weighted clustering coefficient, weighted degree and weighted betweenness) and each column a different social learning rule (left to right: simple, threshold, proportion, conformity). Lines show the predicted effects generated from linear mixed-effect models (LMM) and ribbons show the 95% confidence intervals (see Table S1 for model results). Colour represents the different parameters set for the social transmission rate.

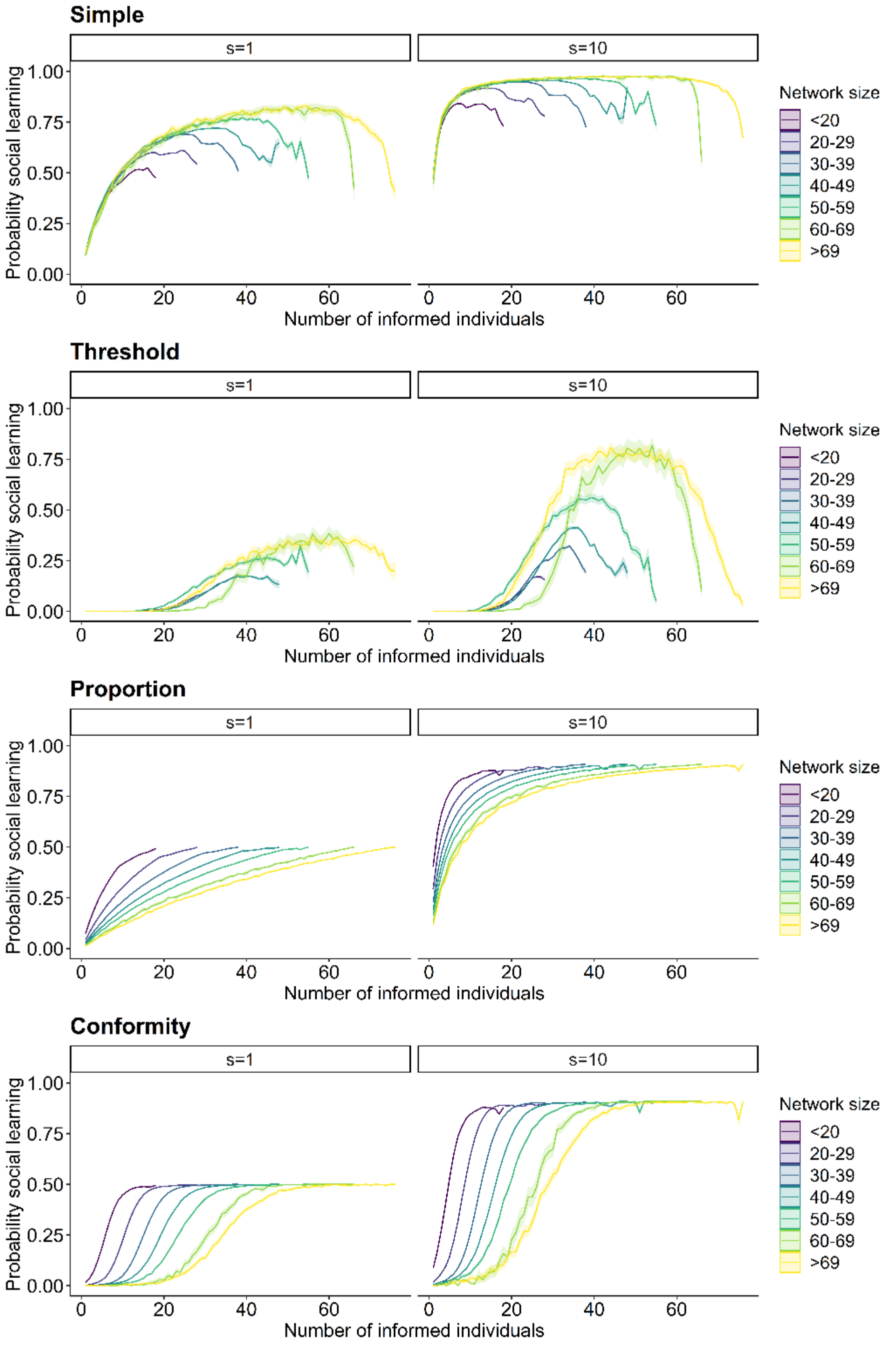


**Figure S14. Relationship between the probability of an individual socially adopting the seeded behaviour and the number of informed individuals within the network under different social transmission rates.** Columns show the results for two different social transmission rates (s=1, s=10, see main text for results on s=5). Rows show results for the four social learning rules (simple, threshold, proportion, conformity). The x-axis describes the number of informed individuals within a social network. At each timestep a new individual adopted the seeded behaviour, whereby each time each individual has a probability of adopting the behaviour through social learning (y-axis) given the set learning rule. Lines plot the average probability for each timestep and ribbons show the 95% confidence intervals from the 100 simulations across the binned groups for different network sizes. Colour represents network size with darker colour indicating smaller networks.

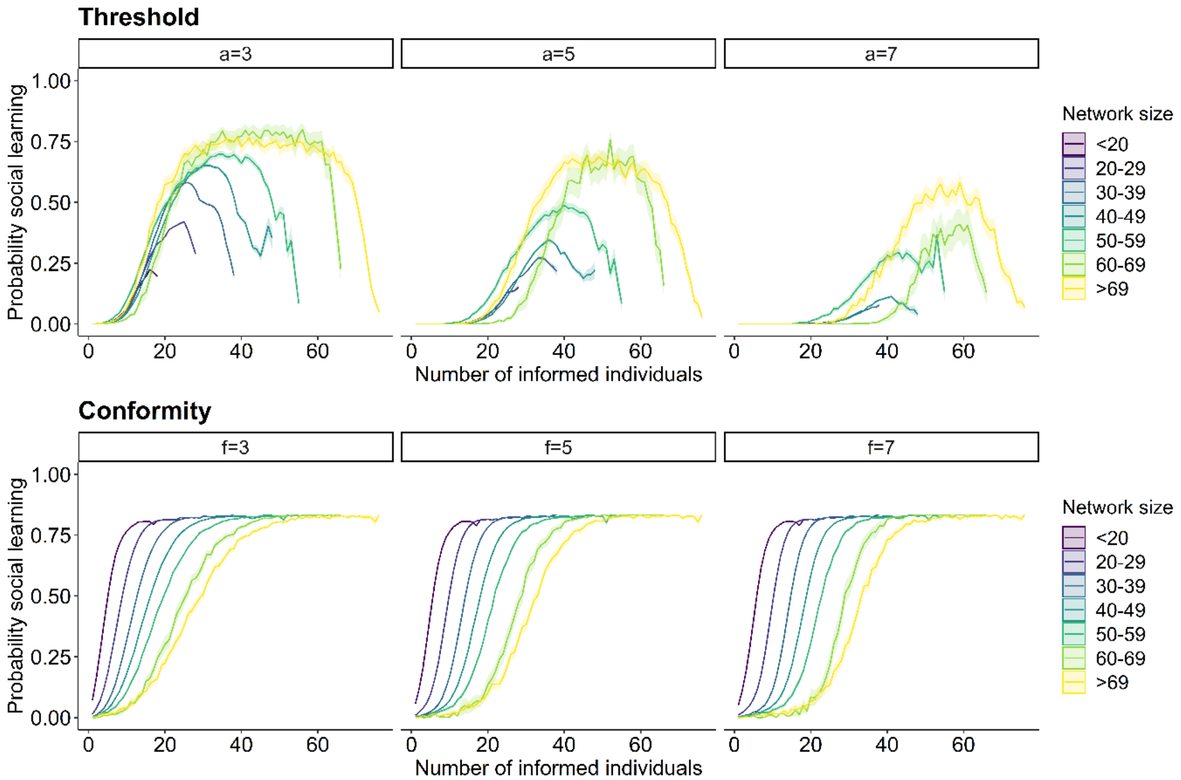


**Figure S15. Relationship between the probability of an individual socially adopting the seeded behaviour and the number of informed individuals within the network under different threshold location (a) and frequency dependence (f) parameters.** Plots on the top show results for the threshold model, plots on the bottom results for the conformity model. Columns show for each model different parameters, i.e. threshold location (a) and frequency dependence (f) parameters (left to right: 3, 5, 7). At each timestep a new individual adopted the seeded behaviour, whereby each time each individual has a probability of adopting the behaviour through social learning (y-axis) given the set learning rule. Lines plot the average probability for each timestep and ribbons show the 95% confidence intervals from the 100 simulations across the binned groups for different network sizes. Colour represents network size with darker colour indicating smaller networks.

**Table S1.** Correlation coefficients between the three individual network metrics.

| | Weighted degree | Weighted clustering coefficient |
|---|---|---|
| **Weighted clustering coefficient** | -0.31 | |
| **Weighted betweenness** | 0.39 | -0.34 |

**Table S2.** Effects of network size on the mean correlation coefficient for weighted clustering coefficient, degree and betweenness for each of the four social learning models.

| Dependent variable | Model | Fixed effects | Estimate | SE | t | P |
|---|---|---|---|---|---|---|
| ***Mean correlation coefficient for weighted clustering coefficient*** | ***Simple*** | Intercept | 0.05 | 0.01 | 3.83 | 0.01 |
| | | Network size | 0.002 | 0.0004 | 6.52 | <0.001 |
| | ***Threshold*** | Intercept | -0.004 | 0.002 | -1.9 | 0.07 |
| | | Network size | 0.0004 | 0.0001 | 4.57 | <0.001 |
| | ***Proportion*** | Intercept | 0.001 | 0.003 | 0.37 | 0.71 |
| | | Network size | 0.0001 | 0.0001 | 1.29 | 0.2 |
| | ***Conformity*** | Intercept | -0.01 | 0.003 | -3.05 | 0.003 |
| | | Network size | 0.0001 | 0.0001 | 1.03 | 0.3 |
| ***Mean correlation coefficient for weighted degree*** | ***Simple*** | Intercept | -0.24 | 0.01 | -20.68 | <0.001 |
| | | Network size | -0.002 | 0.0002 | -8.53 | <0.001 |
| | ***Threshold*** | Intercept | 0.02 | 0.004 | 6.19 | <0.001 |
| | | Network size | -0.002 | 0.0001 | -15.87 | <0.001 |
| | ***Proportion*** | Intercept | -0.06 | 0.005 | -12.54 | <0.001 |
| | | Network size | 0.001 | 0.0001 | 10.06 | <0.001 |
| | ***Conformity*** | Intercept | -0.02 | 0.003 | -8.44 | <0.001 |
| | | Network size | 0.001 | 0.0001 | 7.70 | <0.001 |
| ***Mean correlation coefficient for weighted betweenness*** | ***Simple*** | Intercept | -0.14 | 0.012 | -11.6 | 0.002 |
| | | Network size | -0.001 | 0.0002 | -3.17 | 0.002 |
| | ***Threshold*** | Intercept | 0.01 | 0.002 | 3.62 | <0.001 |
| | | Network size | -0.001 | 0.0001 | -7.32 | <0.001 |
| | ***Proportion*** | Intercept | -0.05 | 0.004 | -12.55 | <0.001 |
| | | Network size | 0.001 | 0.0001 | 9.30 | <0.001 |
| | ***Conformity*** | Intercept | -0.02 | 0.002 | -8.83 | <0.001 |
| | | Network size | 0.001 | 0.0001 | 7.59 | <0.001 |
